## Supplementary Figures for "Genetically encoded transcriptional plasticity underlies stress adaptation in Mycobacterium tuberculosis"

Figure S1.

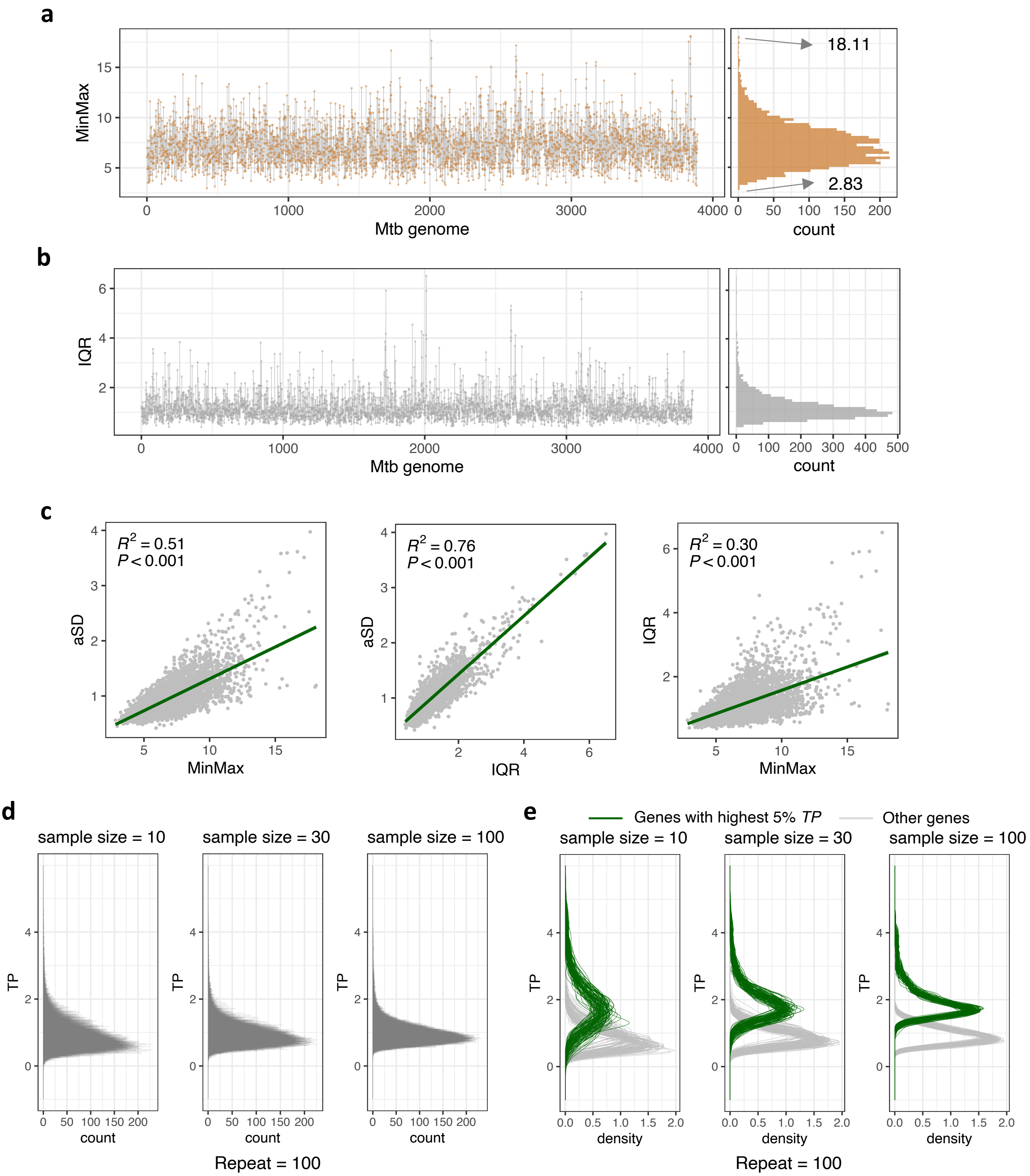


**Supplementary Figure 1 (a-b)** Genome-wide *MinMax* (a) and *IQR* (b) profiles of the 3,891 *Mtb* gene, ordered by their genomic position. The genome-wide distributions of *MinMax* and *IQR* are illustrated in the right panel, with the highest and lowest *MinMax* values being 18.11 and 2.83 respectively, as shown in **(a)**. **(c)** Correlations among adj-*SD* (*aSD*), *MinMax* and *IQR*, with Green lines denoting linear fits. **(d)** Density plots demonstrate the distributions of TP estimated using 10, 30 and 100 randomly selected samples. This bootstrap analysis was performed 100 times for each sampling size, and all repeats are overlaid in each ©t. **(e)** Density plots show the TP distributions of 195 high-TP genes (green) in Fig 2a and the remaining 3,696 genes (grey) at each sample size in the resampling process in Fig. S1d. This bootstrap analysis was repeated 100 times for each sampling size, with all repeats overlapped in each plot.

Figure S2.

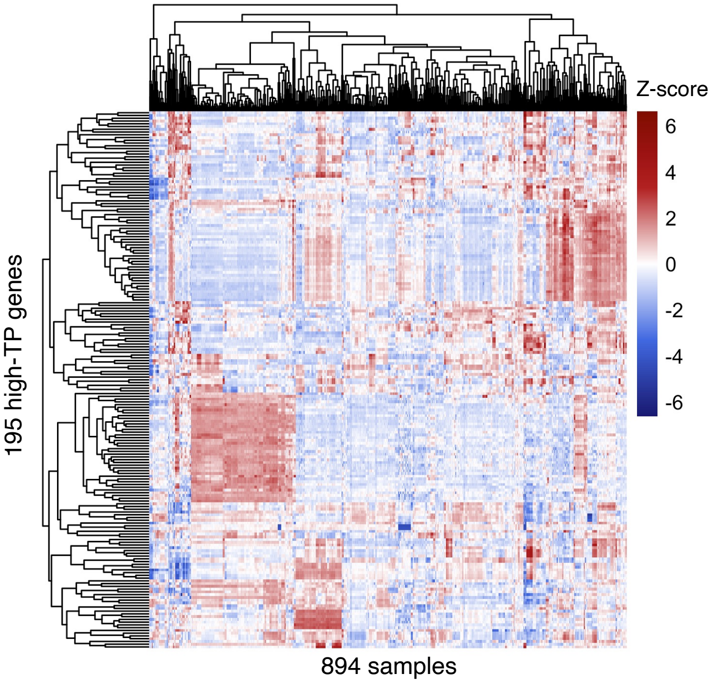


**Supplementary Figure 2** A Heatmap displaying the expression level of 195 high-TP genes across the 894 samples. Expression levels (log RPKM) are scaled and normalized using the Z-score method.

Figure S3.

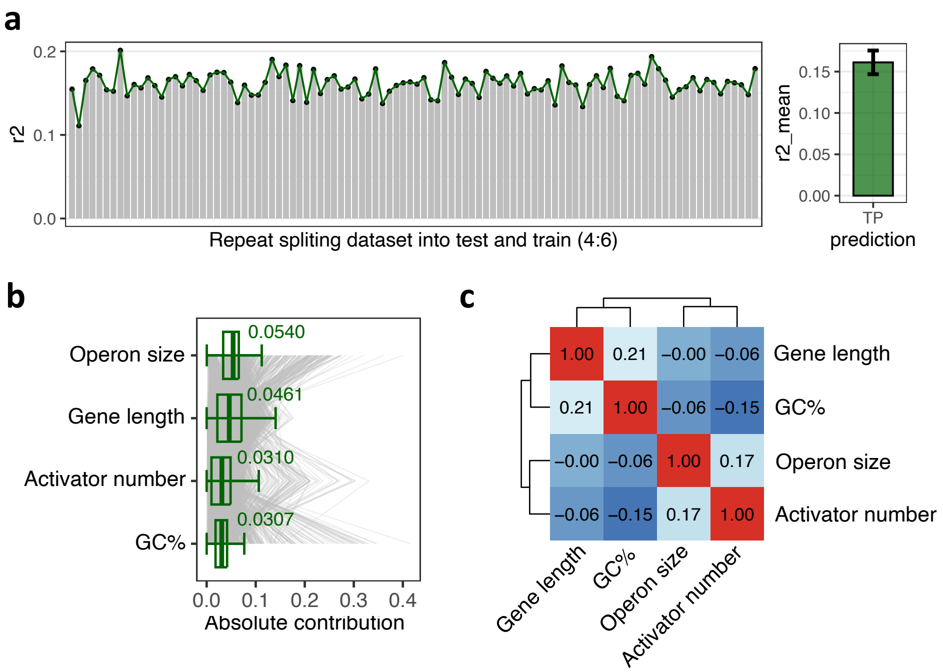


**Supplementary Figure 3 (a)** A barchart depicts the coefficients of determination (R2) of the 100 prediction models described in **Fig. 3b**. Each model was trained on data from 60% of randomly selected genes and the R2 was calculated using the remaining data. The average of the 100 R2 measures are shown in the right panel. The error bar represents the mean ± SD. **(b)** Boxplots illustrate the contribution of the four features to the SVM model described in **Fig 3d**. The contribution of each feature to the SVM model is evaluated using the Shapley additive explanations (SHAP) method, and the absolute SHAP scores were plotted (see *Methods*). Grey lines represent the 2,016 genes involved in the training of the SVM model in **Fig. 3d**. **(c)** A heatmap shows the pairwise Spearman's correlation coefficients among the four impactful genetic features.

Figure S4.

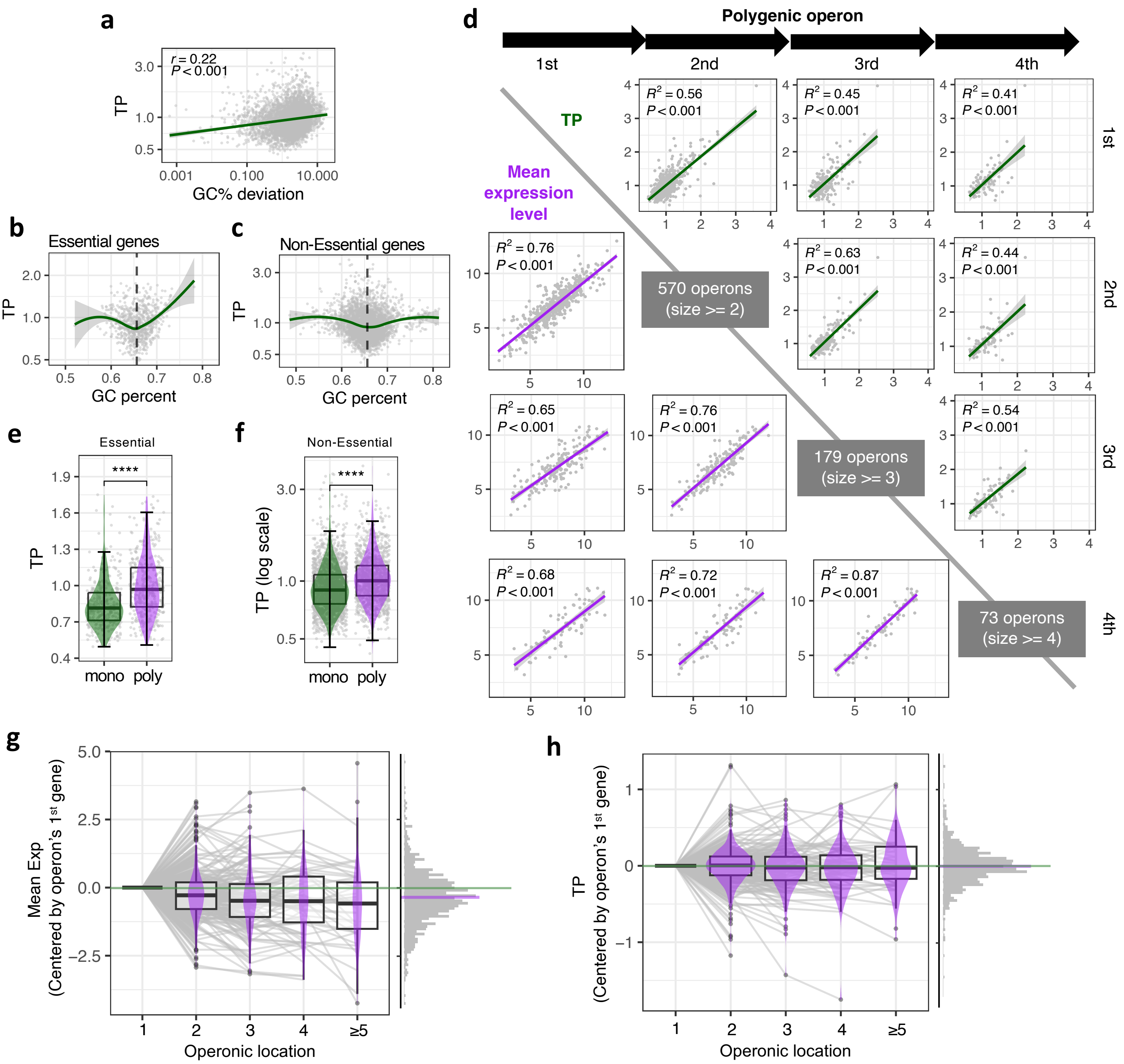


**Supplementary Figure 4 (a)** GC% deviation exhibits a significant positive correlation with TP. The GC% deviation is defined as the absolute difference between a gene's GC% and the genome-wide average. **(b-c)** Scatter plots illustrate the correlations between TP and GC% in essential genes (b) and non-essential genes (c). Green lines represent the LOESS fits, while black dashed lines represent the genome-wide average GC content (65.6%). **(d)** Pair-wise comparisons of TPs and mean expression values among genes located in different positions (1st, 2nd, 3rd, and 4$^{th}$ from the 5' termini of the transcripts, respectively) of polygenic operons. Colored lines represent the linear fits (purple for mean expression level and green for *TP*). **(e-f)** Boxplots demonstrate that both essential (e) and non-essential (f) genes in polygenic operons have significantly higher TP than genes in monogenic operons. Shaded areas depict the TP distribution of monogenic operon genes (green) and polygenic operon genes (purple). Error bars represent mean ± SD. The X-axis in the bottom plot is log-scaled. **** p value < 0.0001. **(g-h)** Boxplots show centered mean expression values (g) and *TP* (h) of each gene in its operon. The X-axis indicates the location of each gene in its operon. Mean expression values or *TPs* of co-operonic genes are centered by the mean expression or *TP* of the first gene in this operon. The normalized mean expression levels or TPs of adjacent operonic genes are connected by grey lines to demonstrate the trend of expression or TP changes along this operon. Density plots represent the aggregated distribution of normalized expression or TP values. Purple lines represent the median values of all genes, and green solid lines represent that of the leading genes (centered to become zero).

Figure S5.

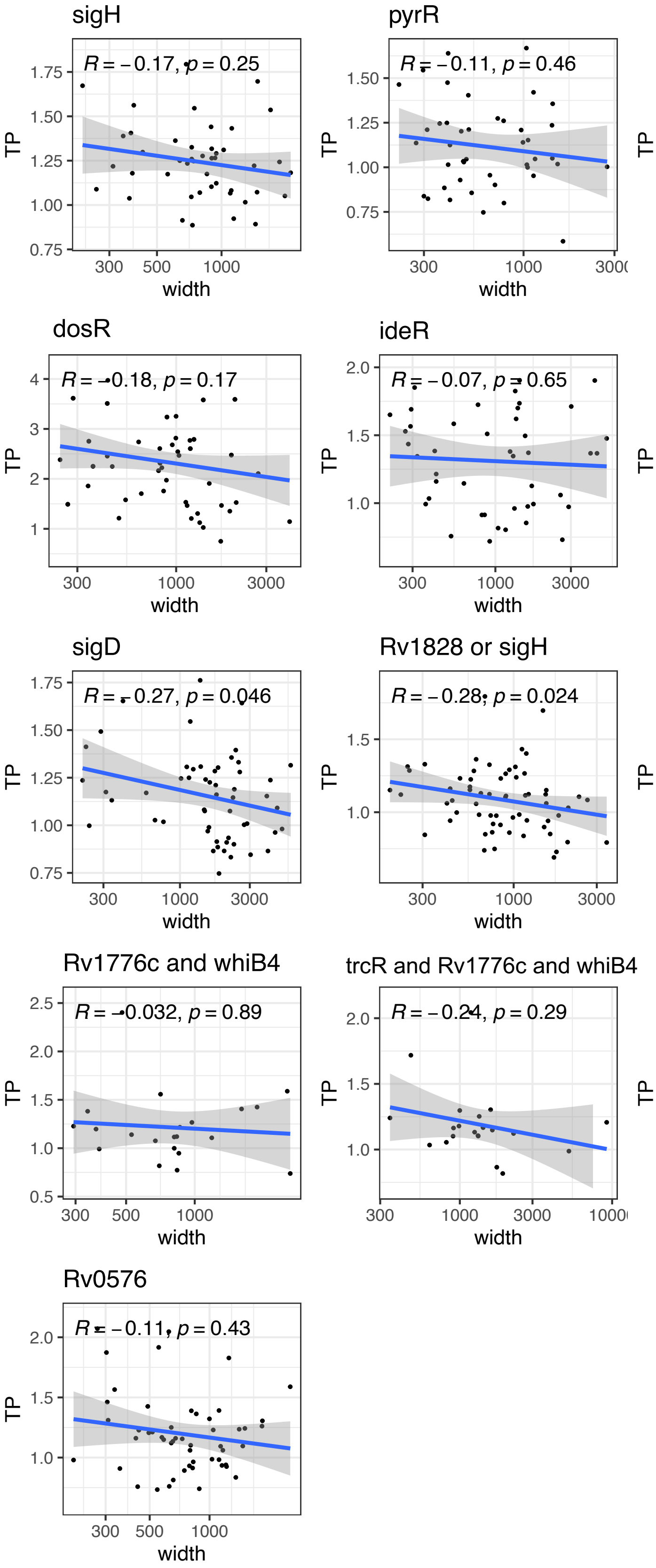


**Supplementary Figure 5** Scatter plots illustrate the correlations between *TP* and gene width in 13 regulons, each containing more than 20 genes ("DevR-1" and "DevR-2" were arbitrarily consolidated into the "DevR" regulon). Lines represent linear fits. Spearman's correction coefficient and corresponding *p* values are provided for each regulon.

Figure S6.

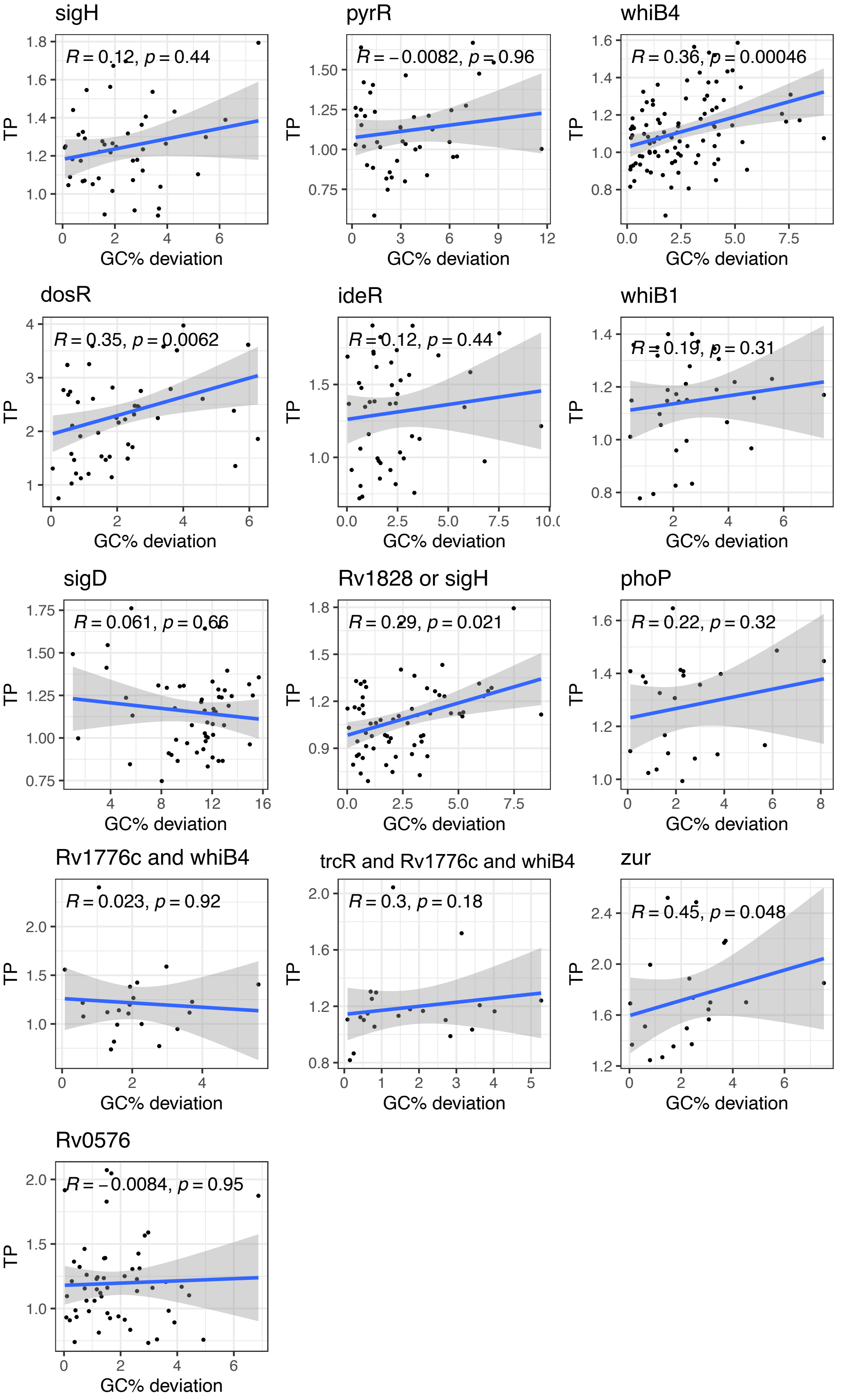


**Supplementary Figure 6** Scatter plots illustrate the correlations between *TP* and GC% deviation in the 13 regulons described in Fig. S5. Lines represent linear fits. Spearman's correction coefficient and corresponding *p* values are provided for each regulon.

Figure S7.

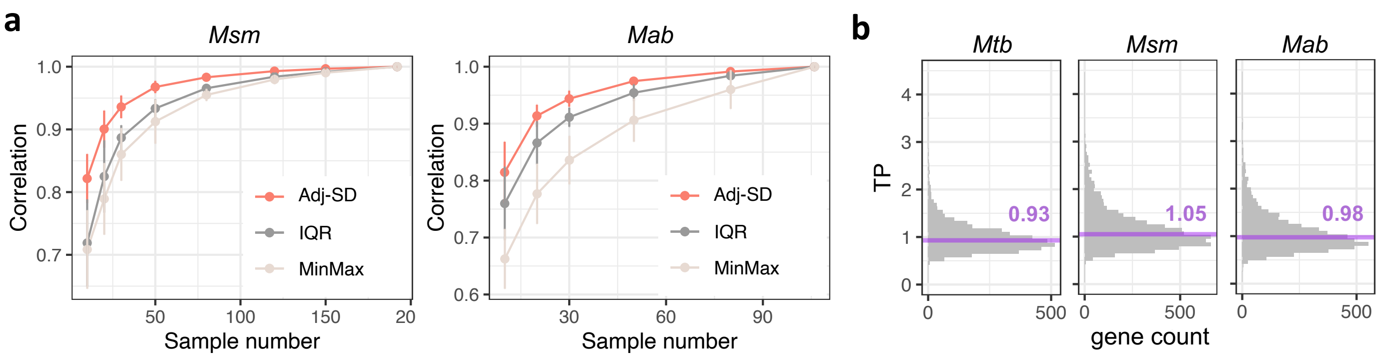


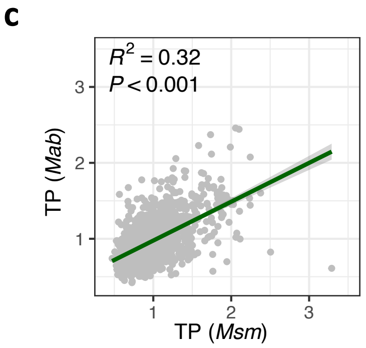


**d**

Functional enrichment of genes with highest 5% TPs

| **Term *(Msm)*** | **FDR** | **Count** |
|---|---|---|
| Universal stress protein A | 0.016 | 6 |
| Iron | 0.063 | 17 |

| **Term *(Mab)*** | **FDR** | **Count** |
|---|---|---|
| Ferritin-like superfamily | 0.024 | 8 |
| Membrane | 0.026 | 43 |

**Supplementary Figure 7 (a)** Comparisons of adj-*SD*, *IQR*, and *MinMax* metrics in describing expression variability of *Msm* and *Mab* genes were made using a bootstrap analysis analogous to the approach described in Fig. 1d. (see ***Materials and Methods***). The lines and the error bars denote the means and the standard deviations of the correlation coefficients rendered by the bootstrap analysis using different metrics. **(b)** *TP* distribution profiles of *Mtb*, *Msm*, and *Mab* genes. Purple lines represent the median TP of all genes for each species. **(c)** Significant correlations of TPs between homologous genes in *Msm* and *Mab* (see ***Materials and Methods***). **(d)** Enrichment analysis conducted on the top 5% highest TP genes in each species (316 genes in *Msm*, 242 genes in *Mab*) and using the DAVID platform. The significance threshold for enrichment results was set at FDR < 0.1.
